# Diet-derived Microbial Metabolites Modulate Stress-Responsive Gene Expression in Germ-free Zebrafish

**DOI:** 10.64898/2026.05.04.722778

**Authors:** Jayson Dale R. Capistrano, Bianka Ketheeswaranathan, Matthew S. Horn, Pham Ngoc Giao Tran, Taylor Ball, Shalvi Chirmade, Sarah J. Vancuren, David W. L. Ma, Kathryn Walton, Emma Allen-Vercoe, Terence J. Van Raay

## Abstract

The gut microbiome plays a pivotal role in overall host health, yet the extent at which diet-derived microbial metabolites affect neurodevelopment and inflammation remains unclear. Here, we employed the *robogut* bioreactor system seeded with fecal samples from two healthy pediatric donors to generate microbial communities exposed to four different diets: low fiber Western (LFW), high fiber Western (HFW), Mediterranean (MED), and Yanomami (YAN), as well as three fiber supplements: fruit and vegetable fiber (FVF), cereal fiber (CRF), and resistant starch fiber (RSF). Metabolites produced by these microbial communities were isolated and applied to germ-free zebrafish (*Danio rerio*) embryos to assess their effects on neurodevelopment and inflammatory gene expression under basal and stress-induced conditions. Despite minimal changes in microbial composition across diets and fiber sources, significant differences in short-chain fatty acid concentrations were observed. Metabolite treatments had limited effects on the expression of neural and inflammatory genes under basal conditions. Under stress conditions, metabolites from any diet mitigated stress-induced *bdnf* expression, suggesting a possible modulatory role of microbial metabolites on stress responses. Overall, these findings highlight the resilience of microbial communities to dietary changes and underscore the importance of microbial metabolite output and its donor-specific nature in influencing host neurodevelopment and immune responses.

## INTRODUCTION

The importance of the gut microbiome in maintaining host health and physiology has been well-established in the literature [1-13]. It is made up of microorganisms that reside within the host gut and encompasses their complete genetic and metabolic functions [14-16]. Its overall composition depends on several factors, both intrinsic and extrinsic, which results in individual hosts harboring unique sets of microbial and metabolic profiles [17-19]. These factors include genetics, geographic location, and diet to name a few. Of these factors, diets have emerged as a strong predictor in determining gut microbiome composition and have been shown to directly impact gut microbial diversity [20-23]. Several studies have shown that consumption of a high-fat Western diet resulted in increased Bacillota/Bacteroidota ratio [24-30] and reduced *Bifidobacterium* spp. [27,30,31], while a prebiotic supplementation resulted in an increase in *Lactobacillus* spp. and other beneficial microbes [2,27,32]. An important function of these gut microbes is to break down indigestible fibers from diets via fermentation [33-36]. In doing so, these microbes produce metabolites which are key effectors of the gut microbiome [32,37]. Interestingly, epidemiological studies have shown that diets play a ubiquitous role in the complex interplay between the gut microbiome and its host [19,31,38-41]. In fact, certain diets have been reported to have beneficial effects, modulate levels of microbial metabolites, and prevent the onset of diseases [31,37-41]. For example, low fiber, high-fat, and high sugar diets have been associated with an increased risk of cardiovascular disease, obesity, and depression [31,42-48], whereas diets rich in fiber and legumes (as in the Mediterranean diet) have been linked to reductions in the onset of chronic disease, anti-inflammatory effects, and better mental health [31,48-52]. Unsurprisingly, perturbations in the gut microbiome are often reported in diseased states [34,53-55], highlighting the involvement of diets in shaping the gut microbiome.

Remarkably, the gut microbiome has been implicated in the development and function of the nervous system [54-60], with many neurological issues having gastrointestinal underpinnings [3,35,61-67]. Predominantly made up of short chain fatty acids (SCFAs) [68], the metabolites produced by the gut microbiome can enter systemic circulation, cross the blood-brain barrier, and ultimately affect the brain [3,35,69-71]. Still, the exact mechanisms and contributions of these metabolites remain elusive. In previous studies, SCFA treatments were reported to alleviate stress, ameliorate depressive behaviors, increase the growth rate of neural cells, and improve memory [1-3,19,27,31,32,35,38,39,69-72]. Although these studies emphasize the neuroprotective effects of metabolites, many of these treatments were limited to individual SCFAs such as sodium butyrate. The gut microbiome, however, produces numerous metabolites [33-35,73,74] which likely work synergistically. Given the role of diets in modulating the gut microbiome and their relationship to both physical and mental health, we endeavoured to understand the effects of metabolites isolated from various diets and evaluate their effects on gene expression using zebrafish as an animal model.

In this study, we employed different diets and fiber sources based on their well-documented effects on the gut microbiome and association with overall health. The low fiber Western (LFW) diet includes foods that are highly processed and contain high amounts of added sugars, fats, and salt. This diet is associated with low consumption of fiber and has been linked to many high-risk diseases and disorders, including obesity, diabetes, anxiety, and depression [31,42-48,75,76]. In contrast, the high fiber Western (HFW) diet follows a largely plant-based diet that often excludes food products of animal origins [77,78], while the Mediterranean (MED) diet is similar but involves the consumption of fish and poultry products [31,48-52,79,80]. Both diets have high fiber contents which is beneficial to the gut microbiome and are linked to better overall health [31,48-52,77-80]. Finally, the Yanomami (YAN) diet is modeled on the substrate consumption of the Yanomami people of the Amazon, many of whom still obtain their sustenance via fishing, gathering, hunting, and simple horticulture [81,82]. Their diets are remarkably low in salt and fat content and are extremely high in fiber [81,82], and those Yanomami who still follow traditional subsistence patterns tend to show very high levels of cardiometabolic health and low incidence of chronic diseases [81-83] which warrants further investigation. We also explored the role of fibers by supplementing soluble and insoluble fibers to the LFW diet due to their known benefits and role in modulating the gut microbiome [83,84]. Soluble fibers are readily metabolized by gut microbes and can therefore, influence the abundance and diversity of the gut microbiota. In contrast, insoluble fibers are poorly metabolized by gut microbes and instead play a role in bulking waste material in the gut, increasing the transit rate throughout the gut and potentially decreasing the time available for gut microbes to ferment non-digested food particles [85]. Given this relationship, evaluating how variations of these fibers affect the gut microbiome is equally important.

## METHODS

### Collection of Fecal Donor Samples

Fecal samples from 10 donors (5 male, 5 female), aged 2 to 6 years old, with a low history of antibiotic use and clinical illnesses and who followed a typical Western diet were evaluated. The samples were collected from participants of the Guelph Family Health Study in accordance with the University of Guelph’s research ethics board approval (REB# 17-07-003) and stored in - 80°C until use. Written informed consent was obtained from each participant or their appropriate guardian.

### Shotgun Metagenomic Sequencing

Fecal DNA was isolated using the QIAamp® PowerFecal® Pro DNA Kit (QIAGEN) and subjected to whole metagenomic shotgun sequencing using Illumina NextSeq 2000 at the University of Pittsburgh’s Microbial Genome Sequencing Center, generating 2x151bp reads with an average of 7,804,105 forward and reverse reads from all samples. Sequencing quality was evaluated using FastQC v.11.9 [89]. Sequences that were less than 50 bp in length were removed using Trimmomatic v.0.39 [90]. Forward and reverse reads were joined using Fastq-join v.1.3.1 [91]. Relative abundance of the microbial taxa was obtained using MetaPhlAn 3 with the ChocoPhlAn 3 database [92]. Bowtie2 was used to align the sequences to clade-specific marker genes [93]. Analyses and visualization on the resulting microbial composition data were carried out in R v.4.1.2.

### Expansion of Fecal Samples

To provide enough biomass for all diet and fiber experiments, the microbial communities from the selected fecal samples were subsequently expanded using bioreactors (Multifors, Switzerland) operated as ‘Roboguts’ under batch conditions for three days [94]. Approximately 3.00g of the selected fecal samples from donors 1 and 2 were used to seed (0.5% w/v) independent bioreactor systems supplied with the HFW diet growth media [94]. Each fecal sample was expanded twice, yielding approximately 1L of biomass, which was divided into 20mL aliquots and stored in -80°C to act as starting inocula for further Robogut experiments.

### Robogut Bioreactor Set-up, Sampling, and Maintenance

The Robogut bioreactor experiment followed the protocol outlined in Gianetto-Hill *et al*. [96]. Growing conditions mimicked the environment of the human distal colon (37°C, pH 7.0, gentle agitation, and under anaerobic conditions from constant sparging with N_2_ gas) and were inoculated with 10% v/v of the expansion culture and allowed to grow in the diet growth media for 24 hours (Day 0). On Day 1, media flow was started with the diet growth media at a rate to match a 24-hour retention time. Overall, both fecal donor samples were exposed to 4 different diets and 3 different fiber sources, with a total of 2 replicates per fecal donors. Formulations of the various diet growth media used are detailed in Table 1 [94,95]. The diet experiments lasted for 21 days and were supplied with either a LFW, HFW, MED, or YAN diet (Figure 1). The LFW diet contains approximately 19g of fiber, while the HFW diet contains 29g. The MED diet has approximately 19g of fiber but also contains legume powder, polyphenols, vegetable peptone, and fish hydrolysate. Finally, the YAN diet contains approximately 33g of fiber and receives a daily supplementation of 1g of fiber from plantain flour and cellulose. For the fiber experiments, the bioreactors were supplied with the LFW diet until day 17 and were subsequently provided with supplements of either fruit and vegetable fiber (FVF; 10g of fiber from pectin), resistant starch (RSF; 9.0g of fiber from corn starch), cereal fiber (CRF; 9.0g of fiber from oat fiber and wheat bran), or left with the LFW diet until Day 28 (Figure 1). The bioreactors were sampled in triplicates on Days 0, 1, 2, then on alternating days and were stored in -80°C. At the conclusion of each experiment, the vessel contents were harvested, aliquoted, and stored in -80°C.

**Figure 1.**
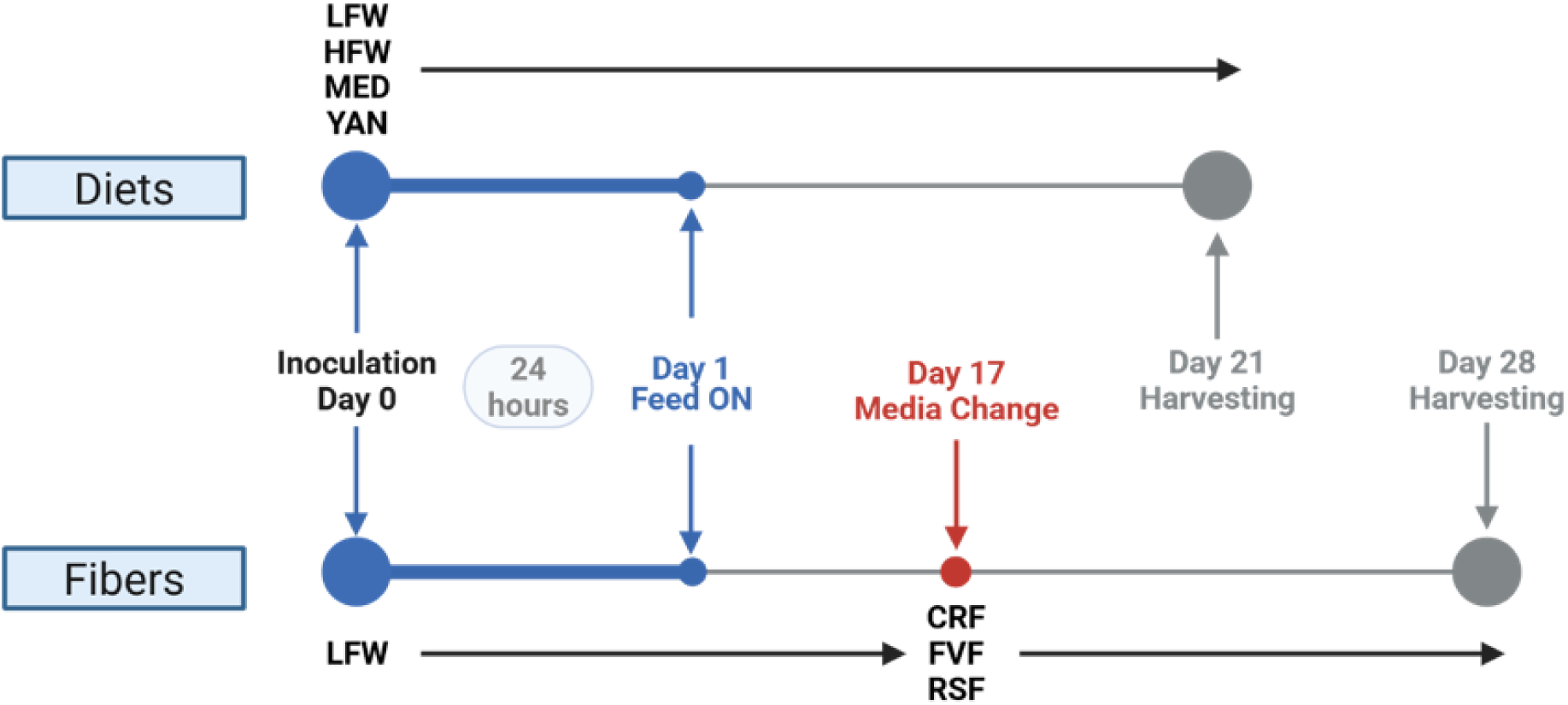
Flowchart of the Robogut bioreactor experiment. All bioreactors were inoculated with the expanded fecal samples on Day 0. Beginning on Day 1, the bioreactors received the diet media at a constant rate. The diet experiments received the same type of diet growth media until Day 21, while the fiber experiments all received the LFW diet until Day 28 or was switched to a different fiber source on Day 17 until Day 28. LFW: low fiber Western; HFW: high fiber Western; MED: Mediterranean; YAN: Yanomami; CRF: cereal fiber; FVF: fruit and vegetable fiber; RSF: resistant starch fiber.

**Table 1.**
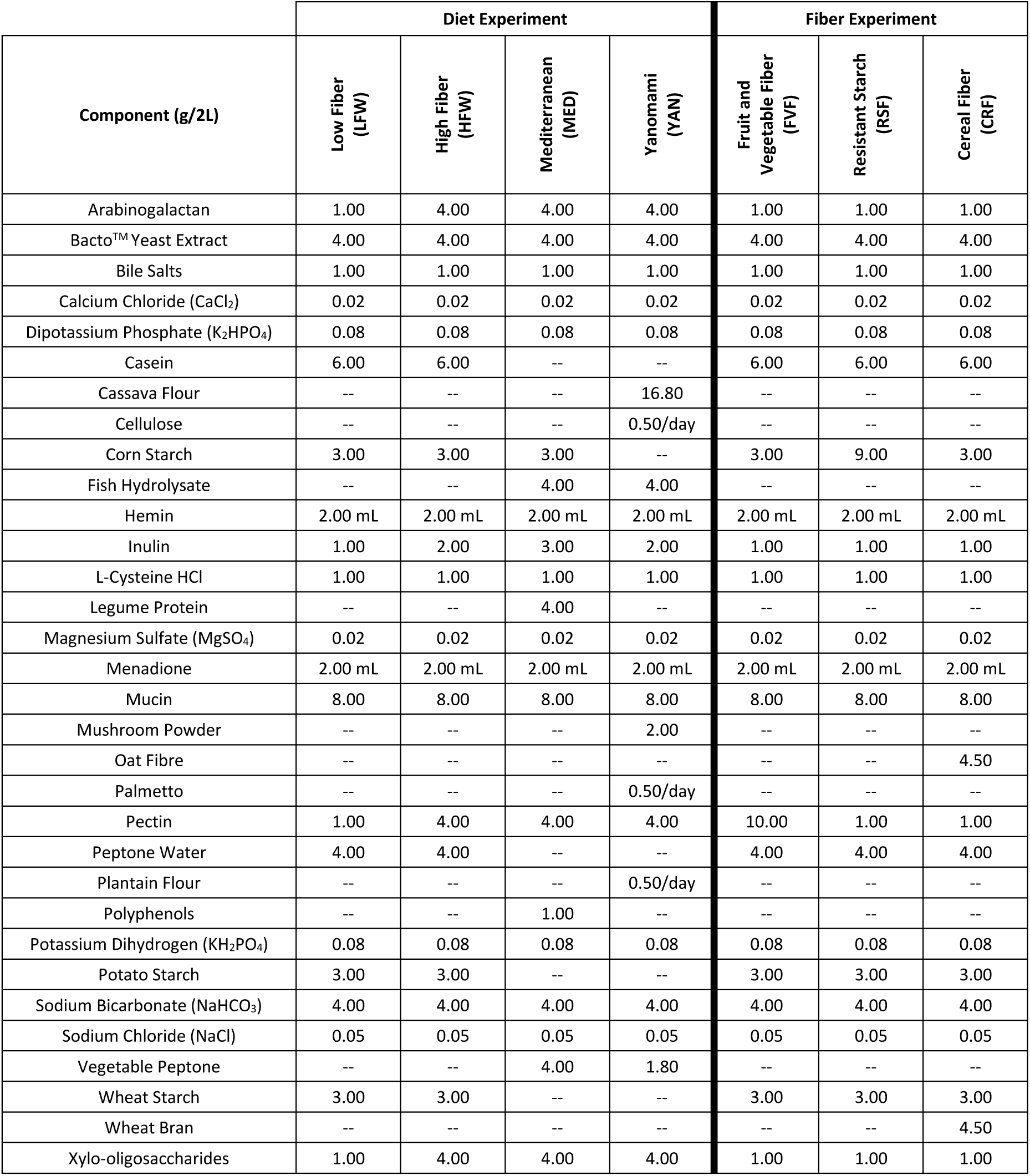
Formulations for each Diet Growth Media.

| Component (g/2L) | Diet Experiment |  |  |  | Fiber Experiment |  |  |
| --- | --- | --- | --- | --- | --- | --- | --- |
|  | Low Fiber (LFW) | High Fiber (HFW) | Mediterranean (MED) | Yanomami (YAN) | Fruit and Vegetable Fiber (FVF) | Resistant Starch (RSF) | Cereal Fiber (CRF) |
| Arabinogalactan | 1.00 | 4.00 | 4.00 | 4.00 | 1.00 | 1.00 | 1.00 |
| Bacto™ Yeast Extract | 4.00 | 4.00 | 4.00 | 4.00 | 4.00 | 4.00 | 4.00 |
| Bile Salts | 1.00 | 1.00 | 1.00 | 1.00 | 1.00 | 1.00 | 1.00 |
| Calcium Chloride (CaCl<sub>2</sub>) | 0.02 | 0.02 | 0.02 | 0.02 | 0.02 | 0.02 | 0.02 |
| Dipotassium Phosphate (K<sub>2</sub>HPO<sub>4</sub>) | 0.08 | 0.08 | 0.08 | 0.08 | 0.08 | 0.08 | 0.08 |
| Casein | 6.00 | 6.00 | -- | -- | 6.00 | 6.00 | 6.00 |
| Cassava Flour | -- | -- | -- | 16.80 | -- | -- | -- |
| Cellulose | -- | -- | -- | 0.50/day | -- | -- | -- |
| Corn Starch | 3.00 | 3.00 | 3.00 | -- | 3.00 | 9.00 | 3.00 |
| Fish Hydrolysate | -- | -- | 4.00 | 4.00 | -- | -- | -- |
| Hemin | 2.00 mL | 2.00 mL | 2.00 mL | 2.00 mL | 2.00 mL | 2.00 mL | 2.00 mL |
| Inulin | 1.00 | 2.00 | 3.00 | 2.00 | 1.00 | 1.00 | 1.00 |
| L-Cysteine HCl | 1.00 | 1.00 | 1.00 | 1.00 | 1.00 | 1.00 | 1.00 |
| Legume Protein | -- | -- | 4.00 | -- | -- | -- | -- |
| Magnesium Sulfate (MgSO<sub>4</sub>) | 0.02 | 0.02 | 0.02 | 0.02 | 0.02 | 0.02 | 0.02 |
| Menadione | 2.00 mL | 2.00 mL | 2.00 mL | 2.00 mL | 2.00 mL | 2.00 mL | 2.00 mL |
| Mucin | 8.00 | 8.00 | 8.00 | 8.00 | 8.00 | 8.00 | 8.00 |
| Mushroom Powder | -- | -- | -- | 2.00 | -- | -- | -- |
| Oat Fibre | -- | -- | -- | -- | -- | -- | 4.50 |
| Palmetto | -- | -- | -- | 0.50/day | -- | -- | -- |
| Pectin | 1.00 | 4.00 | 4.00 | 4.00 | 10.00 | 1.00 | 1.00 |
| Peptone Water | 4.00 | 4.00 | -- | -- | 4.00 | 4.00 | 4.00 |
| Plantain Flour | -- | -- | -- | 0.50/day | -- | -- | -- |
| Polyphenols | -- | -- | 1.00 | -- | -- | -- | -- |
| Potassium Dihydrogen (KH<sub>2</sub>PO<sub>4</sub>) | 0.08 | 0.08 | 0.08 | 0.08 | 0.08 | 0.08 | 0.08 |
| Potato Starch | 3.00 | 3.00 | -- | -- | 3.00 | 3.00 | 3.00 |
| Sodium Bicarbonate (NaHCO<sub>3</sub>) | 4.00 | 4.00 | 4.00 | 4.00 | 4.00 | 4.00 | 4.00 |
| Sodium Chloride (NaCl) | 0.05 | 0.05 | 0.05 | 0.05 | 0.05 | 0.05 | 0.05 |
| Vegetable Peptone | -- | -- | 4.00 | 1.80 | -- | -- | -- |
| Wheat Starch | 3.00 | 3.00 | -- | -- | 3.00 | 3.00 | 3.00 |
| Wheat Bran | -- | -- | -- | -- | -- | -- | 4.50 |
| Xylo-oligosaccharides | 1.00 | 4.00 | 4.00 | 4.00 | 1.00 | 1.00 | 1.00 |

### Extraction of Bioreactor Metabolites

Cell-free metabolites were isolated from the vessel contents of the bioreactor harvested on the last day of each bioreactor experiment (adapted from Ganobis *et al*. [97]). In brief, frozen effluents were thawed at sequentially lower temperatures, first from -80°C to -20°C for at least 12 hours, then -20°C to 4°C for at least 12 hours. Once thawed, the samples were centrifuged at 5,000 × g for 15 minutes at 4°C to remove crude particulate matter, then again at 14,000 × g for 15 minutes at 4°C to remove cells and other remaining particulates. The resulting supernatant was filter-sterilized using a 0.22µM polyether sulfone filter and stored at -20°C until further processing. Every metabolite sample was plated on a non-selective brain-heart infusion (BHI) plate and incubated at 28.5°C for 24 hours to confirm sterility (i.e., zero visible colonies).

### 16S rRNA Gene Sequencing

Genomic DNA was extracted using the QIAmp® PowerFecal® Pro DNA Kit (QIAGEN) following the manufacturer’s instructions. Partial 16S ribosomal RNA (rRNA) gene sequences of the V3-V4 regions were amplified via PCR and sequenced on the Illumina MiSeq platform (2 x 300 bp, at a depth of 100,000 reads per sample) at the University of Guelph’s Advanced Analysis Centre. The R package DADA2 v1.30.0 was used to process paired end reads and create amplicon sequence variant (ASV) tables. Taxonomy was assigned using the *assignTaxonomy* function from the DADA2 package with a set of reference sequences via the Silva 138.1 prokaryotic SSU database. Resulting classified ASV data were imported to Phyloseq v1.38.0 for visualization.

### Nuclear Magnetic Resonance (NMR) Spectroscopy and Analysis

To determine metabolic profiles of the bioreactor samples, prepared metabolites were mixed with 10% Chenomx IS-2 standard containing 4.64mM 3-(Trimethylsilyl)-1-propanesulfonic acid sodium salt in deuterated water and subjected to 1-D ^1^H NMR using the Bruker Avance 600 MHz NMR spectrometer at the University of Guelph NMR Facility. The concentrations of major SCFAs in the NMR spectra were determined via the Chenomx NMR Suite program, version 8.6.

### Zebrafish Maintenance

Adult wild-type Tubingen (TU) zebrafish (*Danio rerio*) lines were raised and maintained in in the Hagen Aqualab at the University of Guelph under approved Animal Utilization Protocol # 4871. Zebrafish were maintained at 28.5°C on a 14:10 light/dark simulated photoperiod and fed commercial flake food or brine shrimp 2-3 times daily.

### Generation and Treatment of Germ-free Zebrafish

Prior to the end of the photoperiod and preceding the day of embryo collection, adult male and female zebrafish were sexed and sorted into spawning baskets with a barricade to separate the sexes. At the onset of the next photoperiod, the barricades are removed to allow natural spawning. Embryos were collected within two hours of fertilization and kept in zebrafish embryo medium (EM; 0.00006% w/v Instant Ocean® Sea Salt solution and 0.001% methylene blue in purified distilled water) at 28.5°C. A clutch of embryos were divided into the various treatment groups and biological replicates were collected from multiple clutches of embryos spawned on different days.

To explore the effects of metabolites, zebrafish embryos were sterilized to remove exogenous bacteria and were made germ-free (GF). Adapted from Pham *et al*. [98] and Rea *et al*. [99], at 30% to 50% epiboly (approximately 5 hours post-fertilization), embryos were divided into conventionally raised (CV) and GF groups. The CV embryos were kept at room temperature (RT), while GF embryos were sterilized at RT to ensure similar development progression. Using sterile technique, the sterilization procedure began with immersing the GF embryos in filter-sterilized gentamicin (100µg/mL) for one hour, followed by 3 five-minute washes in 0.003% filter-sterilized hypochlorite solution and sterile EM performed in a sterile biosafety cabinet. Post-sterilization, the GF embryos were further divided into treatment groups receiving 1% v/v metabolite treatment or no metabolite treatment, with each group having 30-35 embryos each. All embryos were then incubated at 28.5°C. At 1-day post-fertilization (dpf), one embryo from each treatment group was homogenized in a 50uL of sterile EM and a 20µL sample was plated on BHI agar plates, then incubated at 28.5°C for 24 hours to test for sterility. Upon confirmation of sterility (i.e., zero visible colonies), the embryos, as well as the CV embryos, were kept at 28.5°C and harvested at 3dpf.

### Heat, Osmotic, and Swirling Stressor Challenges

At 3dpf, a pool of 10 zebrafish larvae was exposed to 3 different hour-long stressor challenges. For the heat stress, larvae were placed in a 50mL falcon tube containing 20mL EM and submerged in a 37°C water bath (adapted from Lim & Bernier [100]). For the osmotic stress, larvae were transferred into a 10cm petri dish containing 20mL of a 150mM salt solution and incubated at RT (adapted from Cheng *et al*. [101]). Lastly, for the swirling stress, larvae were transferred into a 10cm petri dish containing 20mL EM, placed on a high-speed rotator (Stovall Life Sciences, Inc.), and swirled at approximately 5 rotations per second (adapted from Alsop & Vijayan [102]). Shortly after the stressor, 5 larvae were harvested and macerated in 700uL Trizol, flash frozen at -80°C, and processed for RT-qPCR. A non-stressed, RTcontrol group was included for each stress experiment.

### Quantitative RT-PCR (RT-qPCR)

Total RNA from a pool of five 3dpf larvae was extracted with Trizol following the recommendations in the GENEzol^TM^ TriRNA Pure Kit (Geneaid Biotech Ltd.). RT-qPCR was performed on the CFX96 Touch Real-Time Detection System (Bio-Rad) using the Luna Universal One-Step RT-qPCR Kit (New England Biolabs). Primer efficiencies were measured using SYBR Green Master Mix for qPCR (ThermoFisher) and confirmed to be within 95% to 110% (Table 2) using standard curves. Each biological replicate for RT-qPCR included 3 technical replicates and was normalized to the housekeeping gene, *ef1a*. Gene expressions were calculated using the Livak method and expressed in fold changes.

**Table 2.**
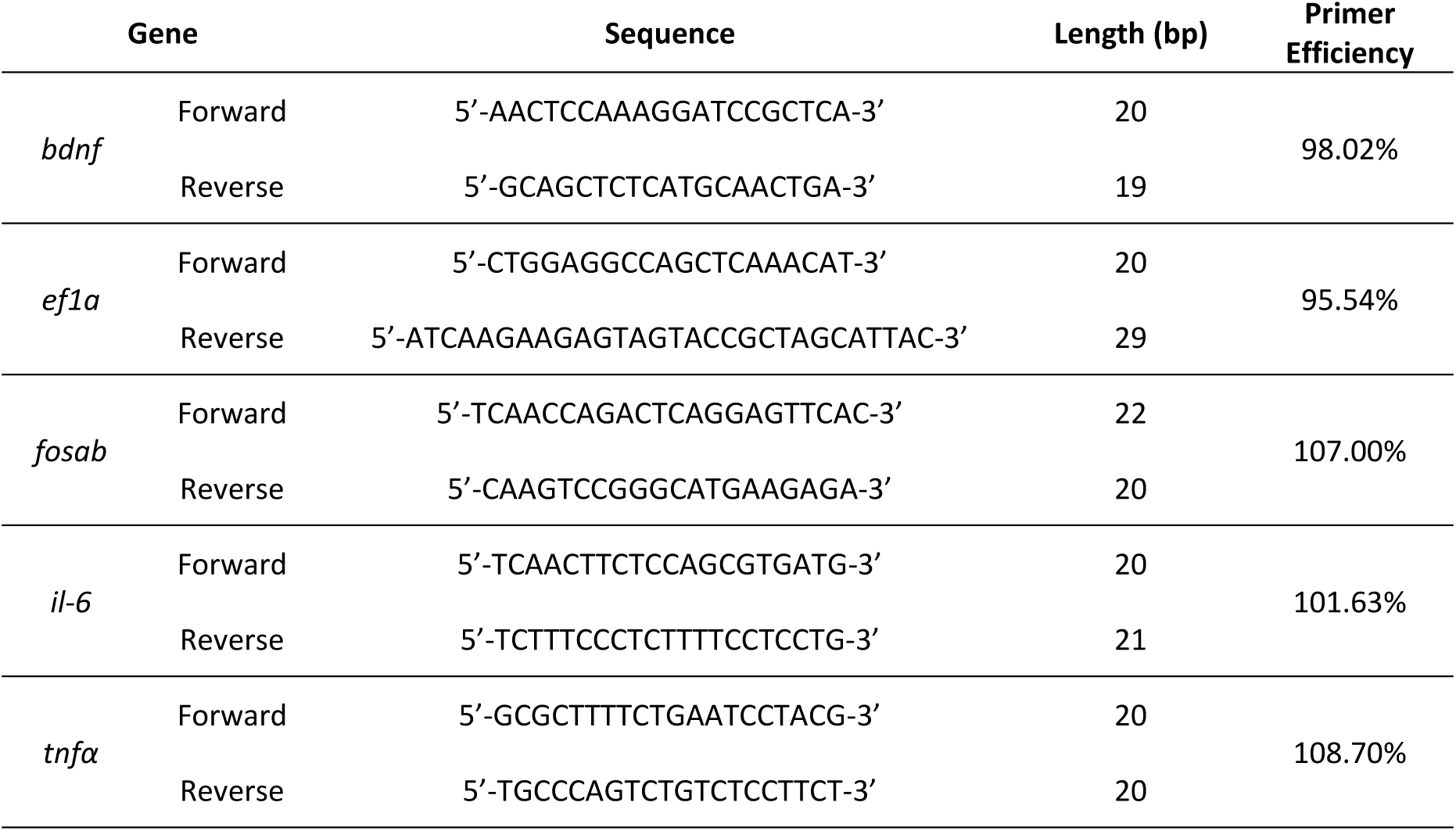
Details of Primer Sequences used in RT-qPCR.

### Statistical Analyses

All data obtained were tested for normality and equal variances. Differences between the treatment groups were analyzed by an analysis of variance (ANOVA) or by a t-test (as specified in each figure) with significance set at p < 0.05. Data sets that do not meet normality and variance assumptions were evaluated using non-parametric tests. All analyses were performed using GraphPad Prism version 10.0.0 for Windows. In the figure legend, uppercase “N” refers to the number of biological replicates.

## RESULTS

### Donor Sample Selection Results

Metagenomic shotgun sequencing of the 10 donor samples was used to determine the taxonomic profiles of each fecal sample (Figure 2A). A Principal Component Analysis (PCA) was used to illustrate the variations between donors so that representative samples could be selected. The PCA plot revealed three clusters, wherein PC1 and PC2 describe 25.83% and 17.88% of the variation, respectively (Figure 2B). A third PC describes an additional 13.60% variation, but no new clusters were formed. In addition, a k-medoid model plot revealed similar findings in that three clusters were formed. In both plots, donors 6 and 8 each formed their own cluster, while the rest of the samples clustered together. From these analyses, donors 1 and 2 were chosen as representatives of the pool as these samples were at the center of its cluster. Donor 1 was a female, 4.3 years old with a history of one antibiotic use (specifically, amoxicillin), and donor 2 was male, 3 years old with two recorded prior antibiotic use (specifically, amoxicillin and ampicillin). The timing of the antibiotic use is unknown.

**Figure 2.**
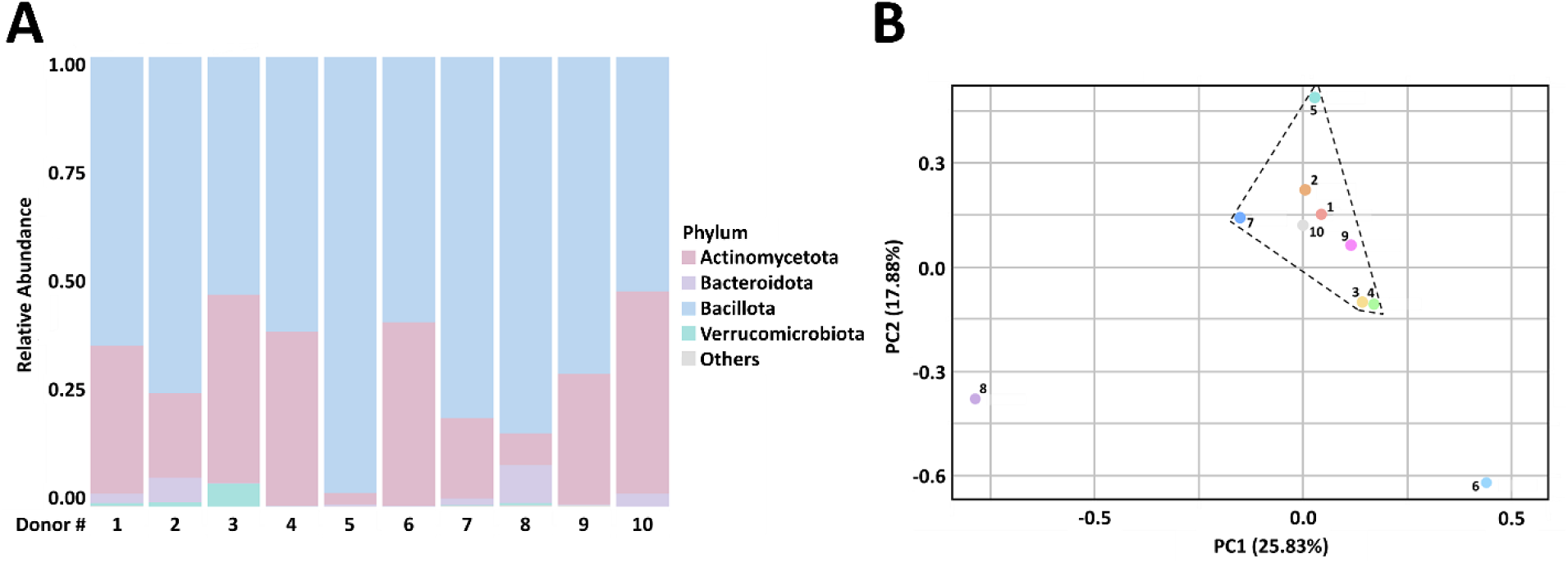
Microbial analysis of ten donor samples. (A) Stacked bar plot showing the top four phyla detected in all 10 donors. The category “Other” describes the phyla with a maximum relative abundance of less than 5% from all samples and was only found in donors 1, 5, 7, 8, and 9. (B) Two-dimensional PCA plot showing the variance between all 10 donors. Enhanced cluster detection using k-medoid identification demonstrated that donors 6 and 8 are separate from the group. From this analysis, donors 1 and 2 were chosen to be the most representative of the group.

### Diets and fiber source did not impact the microbiota cultured in bioreactors

To determine if different diets affected overall microbial profiles, we submitted samples from each diet/fiber source and donor for 16S rRNA gene sequencing analyses at the end of the bioreactor experiments. Despite being supplied different diets, it was revealed that both donors had similar microbial profiles with no clear shifts in microbial compositions or diversity after exposure to each diet (Figure 3A). The distribution of the microbial compositions was found to be consistent between technical replicates, with the most dominant phyla being Bacillota, Bacteroidota, and Verrucomicrobiota. A small portion of the microbes belongs to the Pseudomonadota and Actinomycetota phyla. We then sought to determine if different fiber sources influenced overall microbial profiles but saw no significant changes despite being supplied with different fiber sources (Figure 3B, Supplementary Figure 1). Collectively, this suggests that the different diets and the different fiber sources did not dramatically alter the microbial composition or diversity of the phyla over the course of the experiment.

**Figure 3.**
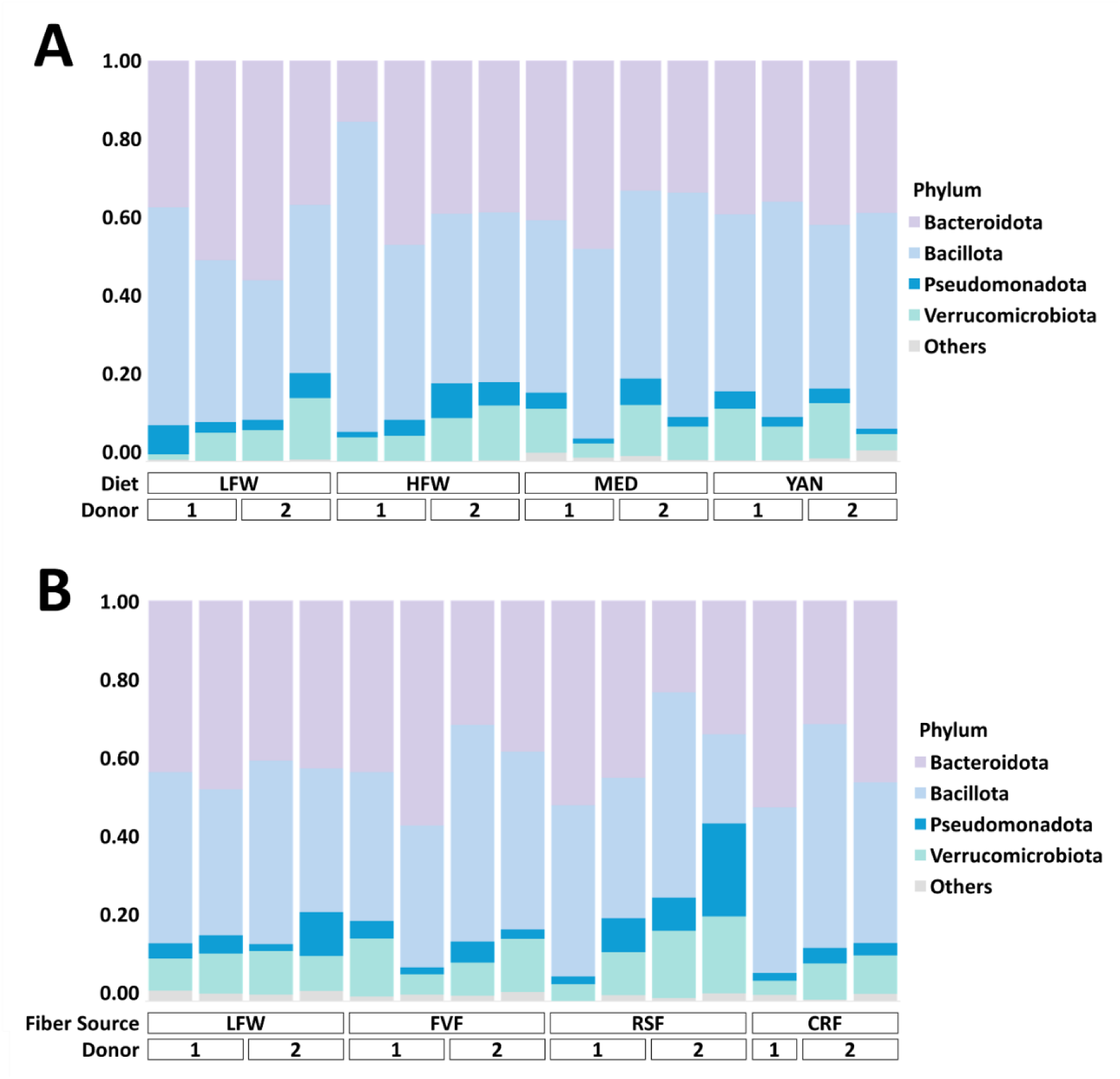
Changes in diets or fiber sources do not significantly alter the microbial composition of the gut microbiome at the phylum level. (A) Bioreactors were supplied with either the LFW, HFW, MED, or YAN diet media for 21 days. (B) Bioreactors were supplied with the FVF, RSF, or CRF diet media at Day 17 and grown for another 10 days; LFW had no added fiber. Effluents were collected at the end of each experiment, and an aliquot was subjected to 16S sequencing. The most dominant phyla found were Bacillota, Bacteroidota, and Verrucomicrobiota. Two biological replicates are shown for each donor, except for CRF for donor 1.

A comparison of the diversity between the raw fecal samples (Figure 2) and the samples collected at the end of both bioreactor experiments (Figure 3) demonstrated that Bacillota remained one of the dominant phyla. Interestingly, we observed a marked decreased in Actinomycetota and an increase in Bacteroidota after both experiments, suggesting that the media used in both experiments may have favored the growth of the latter phyla over the former; however, the microbial phyla seen at the end of both experiments are consistent with the microbes found in the human gut [24,103-109], indicating that the bioreactor is a good model to mimic the environment of the gut.

### Diets and fiber sources impact microbial metabolite output

Despite the lack of appreciable microbial changes between diets or fiber sources, it is still possible that the metabolic profiles between the diets and/or fiber sources are different. To test this, we performed 1-D ^1^H NMR to evaluate changes in the concentrations of the 3 major short chain fatty acids (SCFAs): acetate, butyrate and propionate. There were no significant differences in SCFA concentrations between donors and the changes observed over time were consistent between donors. For the diet experiment, the SCFA concentrations remained relatively stable for the duration of the bioreactor runs (Figure 4A). Here, acetate concentrations were the highest across all diets, followed by propionate then butyrate concentrations; however, in the YAN diet, acetate concentrations were followed by butyrate instead of propionate concentrations. For the fiber experiment, steady increases in SCFA concentrations were observed following the addition of fibers at Day 17 (Figure 4B), with FVF having a significant effect on acetate levels.

**Figure 4.**
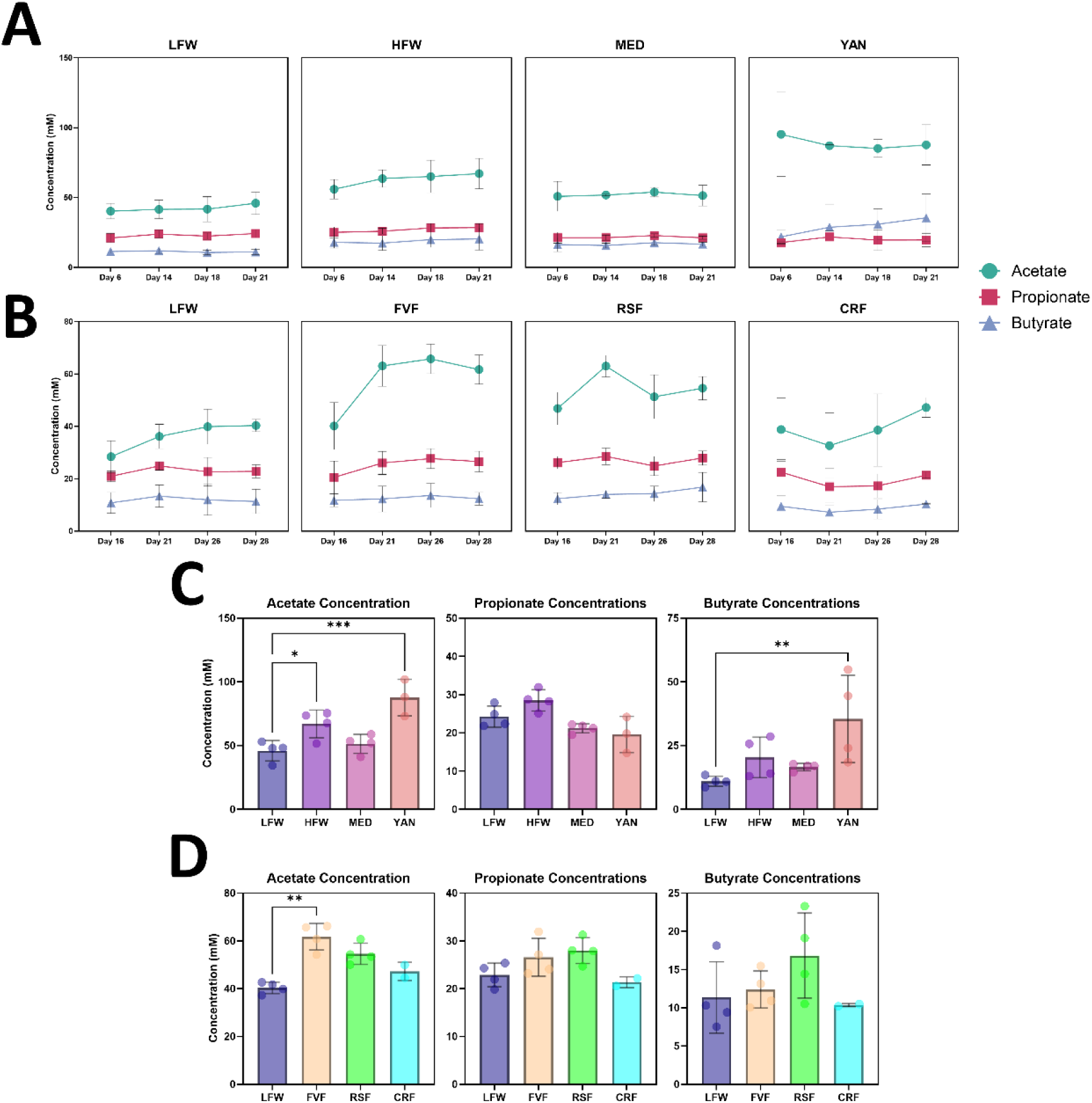
SCFA concentrations reach stability over time. Donors and biological replicates are pooled. (A-B) Metabolites were harvested at the indicated time points and submitted for 1-D ^1^H NMR analysis. In the diet experiment, stability appears to be reached at Day 14, while in the fiber experiment where fiber was added at Day 17, stability appears to be reached at approximately Day 26. In both experiments, acetate concentrations were consistently the highest, followed by propionate then butyrate concentrations (except for YAN). Acetate concentrations were significantly higher on Day 28 when compared to Day 16 in the FVF fiber source (t(6) = 4.16, *p* = 0.025). (C-D) There was a significant effect of diet on acetate (F(3,11) = 11.61, *p* = 0.001) and butyrate (F(3,11) = 10.35, *p* = 0.003) in the diet experiment and a post-hoc test showed that acetate concentrations in HFW (*p* = 0.03) and YAN (*p* = 0.0006) and butyrate concentrations in YAN (*p* = 0.0042) were significantly different from LFW. A significant effect of fiber was found on acetate (F(3,10) = 11.31, *p* = 0.0001), with a post-hoc revealing a significant difference between FVF and LFW acetate concentrations (*p* = 0.004). Error bars represent standard deviation (SD). * *p* < 0.05, ** *p* < 0.01, *** *p* < 0.001 using a one-way ANOVA or Kruskal-Wallis test.

We then compared the SCFA concentrations at the end of each experiment (Day 21 for the diets, Day 28 for the fibers) as these metabolites would be used for the subsequent experiments. Although we could not perform statistical tests for each donor due to the limited number of biological replicates, we did observe some trends (Supplementary Figures 2, 3). For the diet experiment, acetate and butyrate concentrations in both donors tended to be higher in HFW, MED, and YAN when compared to LFW. Propionate concentrations in both donors were highest in HFW but were lower in MED and YAN when compared to LFW (Supplementary Figure 2). For the fiber experiment, all SCFAs saw modest increases in concentrations following the addition of the fibers (Supplementary Figure 3). Overall, both donors had similar metabolite profiles and follows the general 60:20:20 ratio for the 3 major SCFAs. When both donors and replicates are pooled, it was revealed that acetate concentrations from the HFW and YAN diets, as well as the butyrate concentrations from the YAN diet, were significantly different from the LFW diet (Figure 4C). Likewise, the acetate concentrations in the FVF fiber source were found to be significantly higher on Day 28 compared to Day 16 (Figure 4B) but were also significantly different from LFW on Day 28 (Figure 4D). Collectively, these results suggest that diets and fiber impact microbial metabolite output while the microbial composition remains relatively stable.

### The source of metabolites influences gene expression in the developing zebrafish

The gross similarities we observed in the microbial profiles between donors 1 and 2, along with the differences we saw in their metabolic profiles, suggest that subtleties in the microbiome allow individuals to respond differently to the diets and fibers. Given the neuroactive properties of metabolites [1-3,19,27,31,32,35,38,39,69-72], we hypothesized that the different diets and/or fiber sources would ultimately generate metabolites that would affect the nervous system via alterations in gene transcription. Thus, to better understand the contributions of diet on the developing nervous system, we took advantage of the zebrafish model where we previously demonstrated that bacterially derived metabolites are both necessary and sufficient to rescue proper nervous system development [99]. In the current paradigm, we added cell-free metabolites generated from the bioreactors to GF zebrafish embryos at 30 to 50% epiboly. At 3dpf, a pool of five zebrafish larvae was harvested for RT-qPCR to assess changes in the expression of key neural and inflammatory markers (Figure 5).

**Figure 5.**
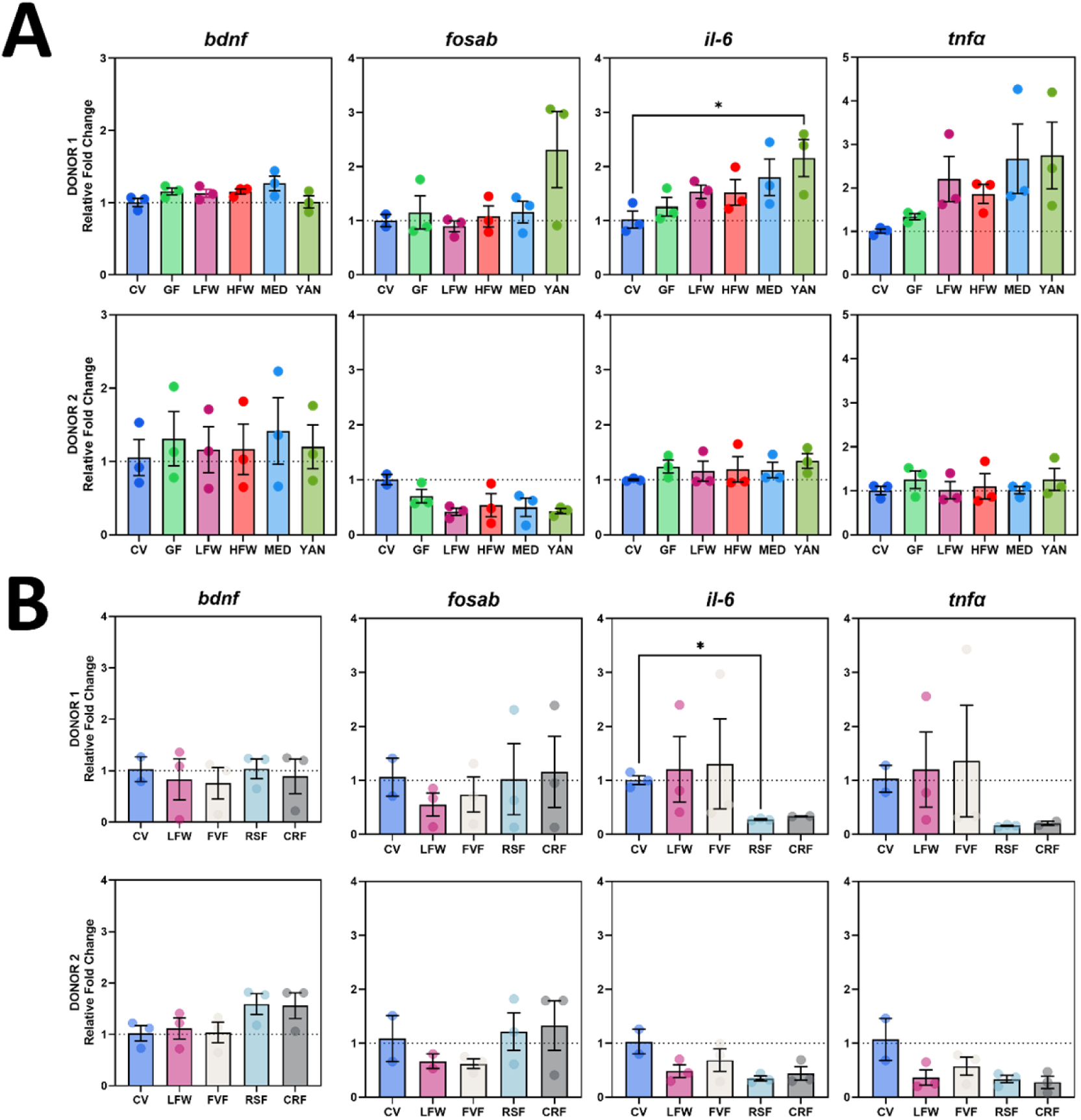
The effects of metabolites on the expression of neurodevelopmental and pro-inflammatory genes were limited, but distinct between donors. GF embryos were treated with metabolites at approximately 5hpf, harvested at 3dpf, and subjected to RT-qPCR. The expression of *bdnf*, *fosab*, *il-6*, and *tnfα* were measured, normalized to *ef1α*, and made relative to the CV group that had received no treatment. GF embryos that did not receive metabolite treatment were also evaluated. (A) Metabolites derived from diets did not seem to affect the expression of *bdnf* and *fosab*. Although no significant effect of metabolites treatment was observed, a post-hoc pairwise comparison revealed that the expression of *il-6* in YAN for donor 1 was significantly different compared to the CV group (*p* = 0.032). (B) Metabolites derived from different fiber source had minimal effects on the expression of the targeted genes, with the exception of *il-6* (F(4,9) = 9.70, *p* = 0.013), noting a significant difference between CV and RSF groups (*p* = 0.034). N = 3. Error bars represent standard error of means (SEM). * *p* < 0.05 using a one-way ANOVA or Kruskal-Wallis test.

Despite the changes observed in the metabolic profiles, this did not translate into significant changes in the expression of the four genes we evaluated; however, we did observe unique expression patterns between donors. When assessing metabolites derived from different diets, the expression of both *il-6* and *tnfα* tended to be higher in zebrafish embryos treated with donor 1 metabolites when compared to CV zebrafish embryos, while *fosab* expression was lower in zebrafish embryos treated with donor 2 metabolites (Figure 5A). For metabolites derived from different fiber sources, we observed that the expression of the four genes evaluated remained relatively consistent between donors, but with a trend of reduced *il-6* and *tnfα* expression in the RSF and CRF fiber sources for both donors. When compared to CV zebrafish embryos, only *il-6* expression in zebrafish embryos treated with RSF metabolites from donor 1 was found to be significantly reduced (Figure 5B). Taken together, these results indicate that the metabolites used in these experiments have limited impact on gene expression in the developing zebrafish nervous and immune systems, at least for the four genes we evaluated.

Given the neuroprotective properties of diets high in fiber, we wanted to determine if the different diets can ameliorate the effects of stressors. We hypothesized that diets rich in fiber (HFW, MED, and YAN) would be able to assuage the effects of stress when compared to the diets low in fiber (LFW). To test this, we chose 3 stressors commonly used in zebrafish studies: swirling, osmotic, and heat [100-102]. First, we evaluated how the stressors affected gene expression in CV and GF zebrafish larvae at 3dpf. For the CV group, the swirling stress caused a significant increase in the expression of *bdnf* (Figure 6A, Controls) and *tnfα* (Supplementary Figure 6A). The osmotic and heat stress significantly increased the expression of *tnfα* only (Supplementary Figures 6B and 6C). We then predicted that the GF group, devoid of microbes and metabolites, would have exacerbated gene expression; however, we only observed significant increases in *bdnf* (Figure 6A, Controls) and *fosab* expression (Supplementary Figure 4A) for the swirling stress, while we saw significant increases in *bdnf* (Figure 6B, Controls) and *fosab* expression (Supplementary Figure 4C) for the osmotic and heat stress, respectively. There were no significant changes in *il-6* expression for CV or GF under any of the stressors (Supplementary Figure 5). These results indicate that the expression of *bdnf* seems to be the most sensitive to the three stressors, with the swirling stressor having a greater effect than the other two stressors.

**Figure 6.**
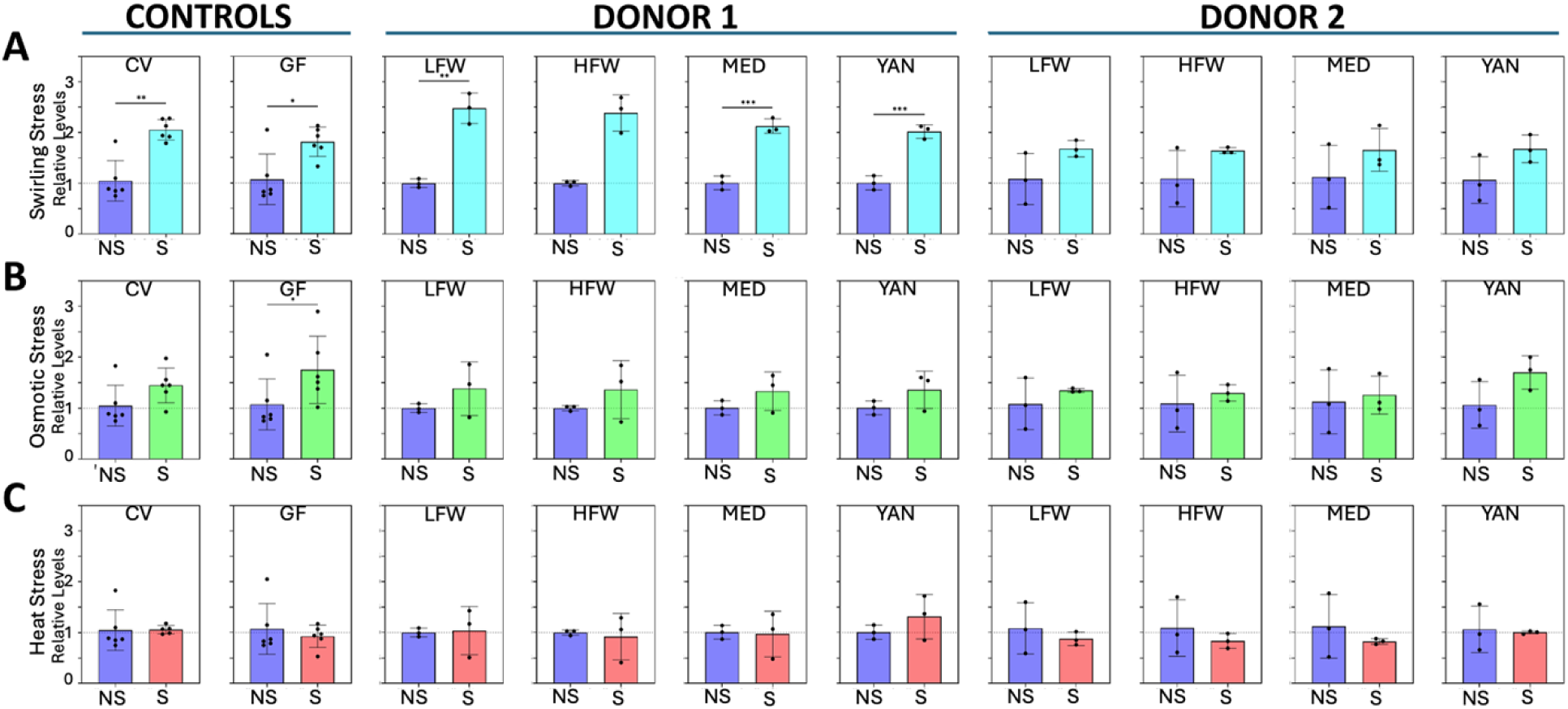
The metabolites can ameliorate the effects of stressors on *bdnf* expression. (A) A significant increase in *bdnf* expression was observed following an hour-long swirling stressor in the CV and GF groups. Similar findings were seen in the LFW, MED, and YAN groups for donor 1, but these were reduced to non-significant levels for donor 2. (B) Only the GF group saw a significant increase in *bdnf* expression following the osmotic stressor. The CV and all metabolite-treated groups were non-significant. (C) The heat stressor did not affect the expression of *bdnf* in all groups. N = 6 for controls, N = 3 for metabolite treatment groups. Error bars represent standard deviation * *p* < 0.05, ** *p* < 0.01, *** *p* < 0.001 using an unpaired t-test or Mann-Whitney test.

Next, we added the different metabolites to GF embryos and evaluated gene expression under the same stressors. While we observed a significant effect of diets, contrary to our predictions, we did not observe any diet-specific effects and the LFW diet affected gene expression similar to the other diets (Figure 6). More specifically, we observed that *bdnf* expression was rescued (i.e., reduced to non-significant levels) by metabolites derived from donor 2, but not donor 1 in the swirling stress (Figure 6A). In contrast, metabolites from both donors were able to rescue *bdnf* expression from the effects of osmotic stress (Figure 6B). Curiously, the heat stress did not seem to impact *bdnf* expression (Figure 6C). After evaluating the expression of *fosab*, *il-6*, and *tnfα*, it was revealed that *fosab* had variable and inconsistent results (Supplementary Figure 4), while both *il-6* (Supplementary Figure 5) and *tnfα* (Supplementary Figure 6) seemed to be relatively stable following all three stressors. Overall, no discernable differences between diets could be identified. Nonetheless, we saw that *bdnf* expression appears to be amenable to both stressor and metabolites and likewise, follows a donor-specific pattern of expression consistent with our previous results.

## DISCUSSION

The role of the gut microbiome in maintaining overall host health has been well-documented in the literature [1-13], but recent studies have also revealed its importance in the function and development of the nervous system [54-60]. Its key effectors, microbial metabolites, have gained increasing attention for their ability to influence the brain and behavior [1-3,19,27,31,32,35,38,39,69-72]—further emphasizing the involvement of the gut microbiome in the nervous system. Previous work in our lab demonstrated that metabolites are necessary for timely neurodevelopment. In the absence of microbes and their metabolites, broad neural gene expression was delayed but was partially rescued by the addition of gut-derived zebrafish metabolites [99]. Predicated on this work and given the documented neuroactive properties of metabolites, we employed 4 different diets (3 of which have been well-studied: LFW, HFW, MED and one unknown: YAN) and 3 different fiber sources (pectin from FVF; wheat bran and oat fiber for CRF; resistant corn starch for RSF) and evaluated how they impacted the microbial and metabolic profiles of two independent gut microbiomes derived from young children with limited antibiotic exposure. We then sought to determine the effects of the metabolites derived from these diets and fiber source on neurodevelopment and inflammation using zebrafish as an animal model.

Although the core microbial composition of the gut microbiome is relatively resilient to disruptions, several studies have reported that these microbes vary daily and can change immediately following a treatment and/or perturbation [110-123]. In fact, there are clinical studies demonstrating changes in microbial composition as early as a few hours following a change in diet, antibiotic treatment, or pre/probiotic supplementation [5,110-123]. Collectively, these studies indicate that the microbes of the gut microbiome are robust and sensitive; however, other studies have described a higher level of resilience in some gut microbiomes. In these studies, dietary interventions did not cause changes in the gut microbiomes even after prolonged exposure [4-6,124-127]. These individuals were deemed ‘non-responders’. In the present study, a decrease in Actinomycetota and an increase in Bacteroidota phyla were seen after the raw fecal samples were expanded. During the bioreactor experiments, no apparent changes or differences were noted in the microbial compositions at the phylum level of our selected donors despite continuous exposure to a consistent diet or the addition of fibers; hence, it is possible that the expansion of the fecal samples made the microbial communities less responsive to the media changes or that donors selected for this study are non-responders as well. Similarly, the gut microbiome of healthy individuals is typically characterized by its great resilience to perturbations [128-131], suggesting that the use of healthy donor samples may explain the lack of significant changes we observed. Because our study was limited to only two healthy donors, future iterations of this experiment should expand the number of donor fecal samples used to increase both the sample size and statistical power, as well as include samples from diseased or unhealthy donors. Indeed, preliminary data from our lab has shown that GF zebrafish embryos treated with metabolites from either a neurotypical or autistic donor have markedly different phenotypes.

Despite the lack of microbial distinctions, we found significant differences in the metabolic profiles between the diets and fiber sources in a pooled analysis. Indeed, significant differences in SCFA levels between LFW and the other diets and fiber sources were found, indicating that high fiber diets or the supplementation of fiber can alter the metabolic profiles of the gut microbiome. Combined with the sequencing results, the differences in SCFA concentrations are strikingly similar to the observations reported by Wu and colleagues [6]. In their study, they report that healthy vegans and omnivores had similar gut microbial profiles. While they did not see significant differences in fecal SCFA concentrations after 10 days of dietary intervention, they attributed this to a high intersubject variability of baseline SCFA concentrations, suggesting that SCFA concentrations depend on the subject, consistent with the donor effects observed in our study. Further, work by Fragiadakis and colleagues [132] revealed that dietary interventions caused substantial changes in participant gut microbiomes, but their microbiomes eventually returned near its baseline status despite maintaining the dietary interventions. Together, these studies corroborate our results, underscoring the resilience of the gut microbiome despite what is typically reported.

In our study, we evaluated the expression of neural markers: brain-derived neurotrophic factor (*bdnf*), *fosab* (a zebrafish homolog of mammalian *fos*), and inflammatory markers: interleukin-6 (*il-6*) and tumor necrosis factor alpha (*tnfα*). Both *bdnf* and *fosab* are involved in neuronal cell differentiation, growth, proliferation, and survival [133-142], making them excellent markers for neurodevelopment. Meanwhile, the innate immune system in zebrafish is functional during early developmental stages [143,144] and serve as its first line of defense against infections [143-146]. In GF animals, the immune system is reported to be blunted and generally have lower levels of circulating cytokines such as IL-6 and TNFα [9,147-153]. However, in our study we did not observe decreases in *il-6* or *tnfα* expression in GF embryos. Furthermore, our results demonstrated that overall, metabolites have limited effect on the expression of the four genes evaluated but demonstrated some donor specific effects, especially in the expression of *il-6* and *tnfα*. These results suggest that the effects of the metabolites depend minimally on the diet or fiber source but are more influenced by its source microbiomes (i.e., the donors).

There are numerous studies demonstrating that SCFA treatment can alleviate stress [1-3,19,27,31,32,35,38,39,69-72], indicating that the effect of diets may only manifest under stressful conditions. As such, we subjected a pool of zebrafish larvae to one of three stressors and found that specific stressors (e.g., osmotic and swirling) evoked increases in expression of our target genes. For example, we saw a significant increase in *bdnf* expression in CV and GF larvae following an osmotic or swirling stress. When treated with metabolites, we found that the four diets were successful in rescuing *bdnf* expression in both donors following an osmotic stressor, but this rescue was only seen in larvae treated with donor 2 metabolites following a swirling stressor. Curiously, the same patterns are not seen in the other genes evaluated. A study by Pate and colleagues [154] found reductions of baseline *fos* in the prefrontal cortex of GF animals, whereas Yu and colleagues [155] saw an increase in *fos* expression in the amygdala and hypothalamus of GF mice. Therefore, the variability of *fosab* expression in our experiments may be due to the use of the whole embryo instead of just the brain or specific brain regions.

Although our results did not support our hypothesis that diets high in fiber would ameliorate the effects of stress, it did demonstrate that metabolites can potentially limit the effects of stress and that its source plays a larger role in determining its effects. Other studies investigating the relationship of the gut microbiome and development often use commercially available, purified, and individual SCFAs and have seen significant effects [2,21,32]. However, the composition of metabolites produced by the gut microbiome is a mixture of many SCFAs and other compounds [33,34,35,73,74] and are likely more dynamic, extending beyond the predominant acetate, propionate, and butyrate compounds. Our study overcomes this limitation by utilizing metabolites directly produced by microbes cultured from our selected donor fecal samples and thus, provides a more physiologically relevant representation of the by-products of these microbes and their effects on a developing system. The results of our study showed that while the microbial component of the gut microbiome is resilient to changes in diet or fiber source, their metabolic output is not, arguing that it is more likely the collective genetic complement of the gut microbiome that matters and not the individual microbial species. Despite the lack of distinctions between the different diets or fiber sources, our study has demonstrated that metabolites can modulate gene expression in response to specific stressors and that these effects depend on the source of the metabolites.

## DATA AVAILABILITY STATEMENT

The data that support the findings of this study are available from the corresponding author upon reasonable request. The data analyzed during this study are publicly available in the NCBI BIOProject accession number PRJNA1391537.

## ETHICS and ANIMAL APPROVAL

The samples were collected from participants of the Guelph Family Health Study in accordance with the University of Guelph’s research ethics board approval (REB# 17-07-003). The University of Guelph’s Animal Care Committee has approved the care and use of the animals for this study (Animal User Protocol #4871).

## CONTRIBUTIONS

TJVR and EAV conceptualized the study; JDRC, BK, TB, MH, and SC completed the data analysis; JDRC, BK, TB, PNGT, MH, SJV performed the experiments; DLM, KW provided access to the GFHS; JDRC, BK, TJVR wrote the manuscript; TJVR oversaw all aspects of the study. All authors have read and approved the manuscript.

## ACKNOWLEDGEMENTS

We would like to thank Hagan AquaLab staff Matt Cornish and Mike Davies for their assistance with our zebrafish colony. Thanks to members of the Van Raay lab and thanks to members of the Allen-Vercoe lab for their assistance with everything microbial. The authors acknowledge the families who participated in the Guelph Family Health Study, Angela Annis and Madeline Nixon for their work in coordinating the GFHS, and Adam Sadowski and Amar Laila for their data analytical expertise.

## List of Figure Legends

**Supplementary Figure 1.**
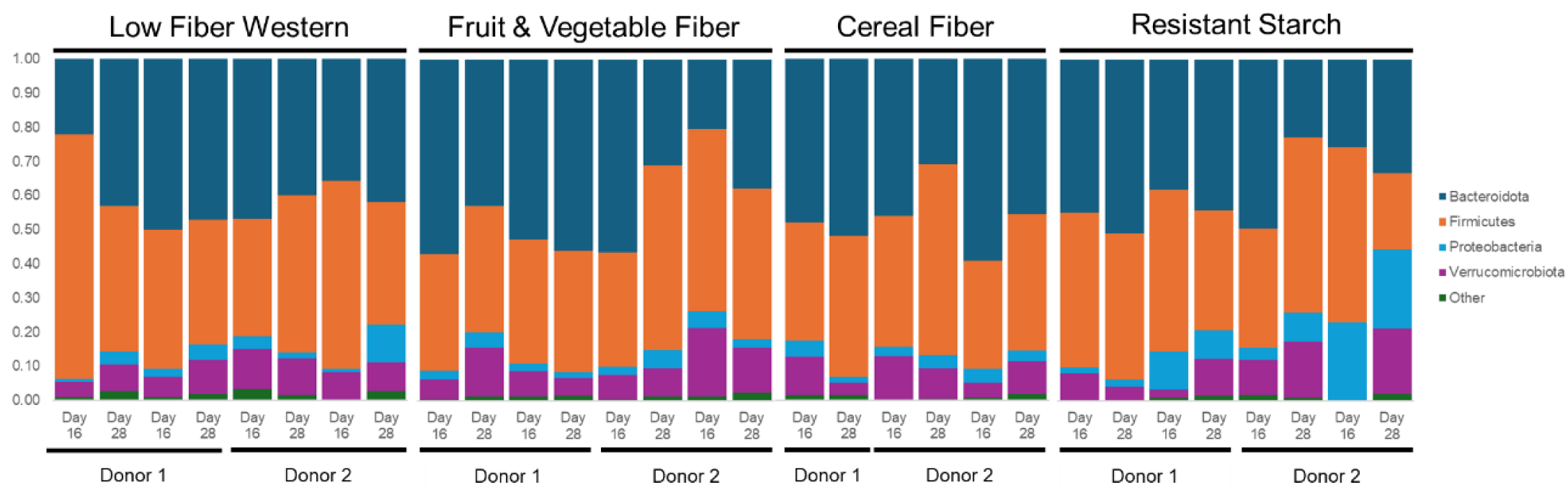
Comparison of the microbial composition at the phylum level before (Day 16) and 10 days after fiber additions at day 28. In the fiber experiment, media was changed and supplemented with different fiber source on Day 16. Despite the addition of fiber for approximately 10 days, no major changes in the microbial composition at the phylum level were noted. Two biological repetitions (Rep) for each donor except for cereal fiber for which there is only 1 rep for donor 1.

**Supplementary Figure 2.**
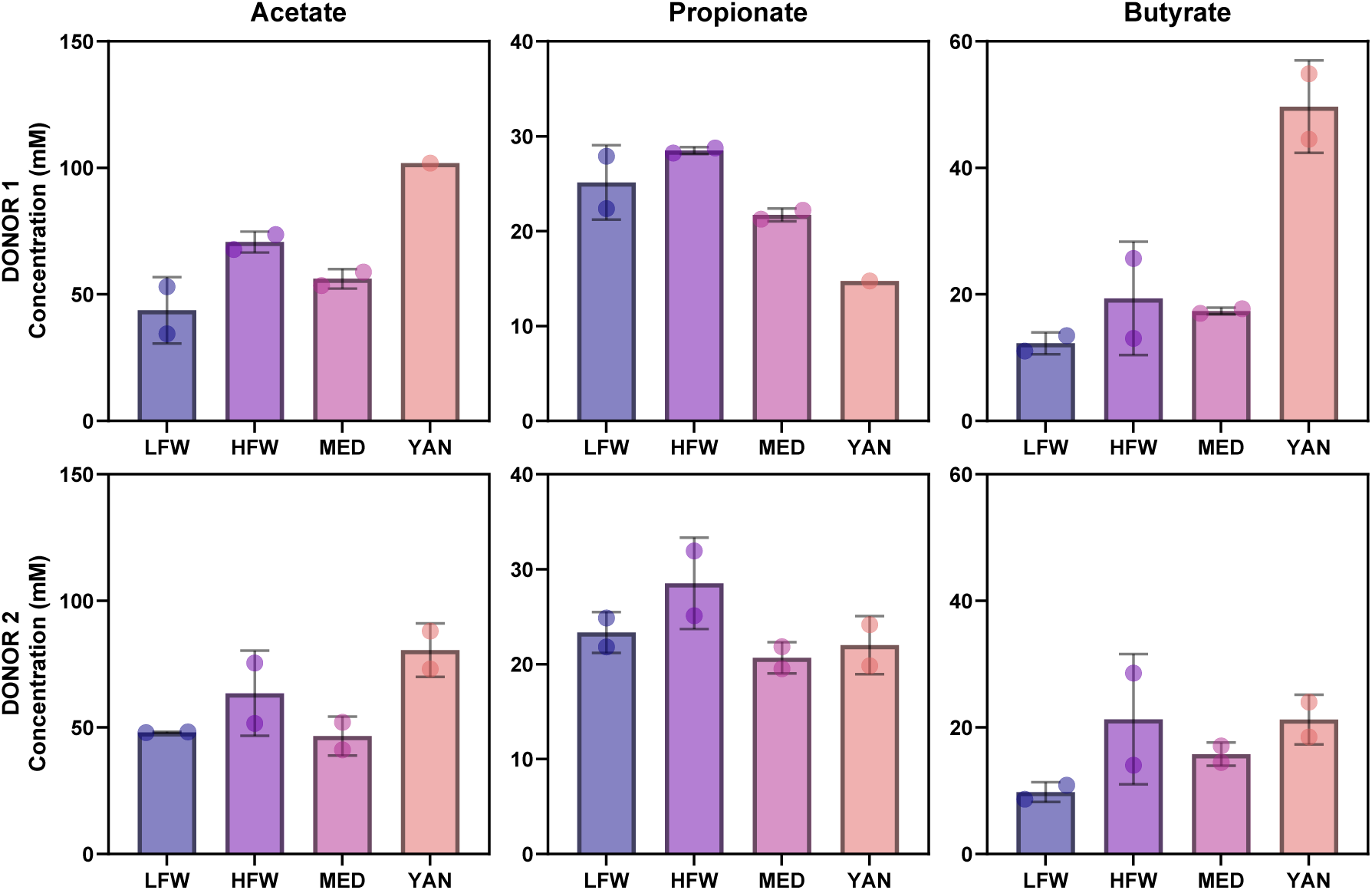
Comparison of Donor 1 and Donor 2 SCFA Concentrations at Day 21 of the Diet Experiment. In both donors, the YAN diet produced the highest average concentrations of acetate while the HFW diet produced the highest average concentrations of propionate. Except for the YAN diet, acetate concentrations were found to be the highest followed by propionate and butyrate concentrations. In donor 1, acetate concentrations were the highest but is followed by butyrate, then propionate concentrations, setting it apart from donor 2.

**Supplementary Figure 3.**
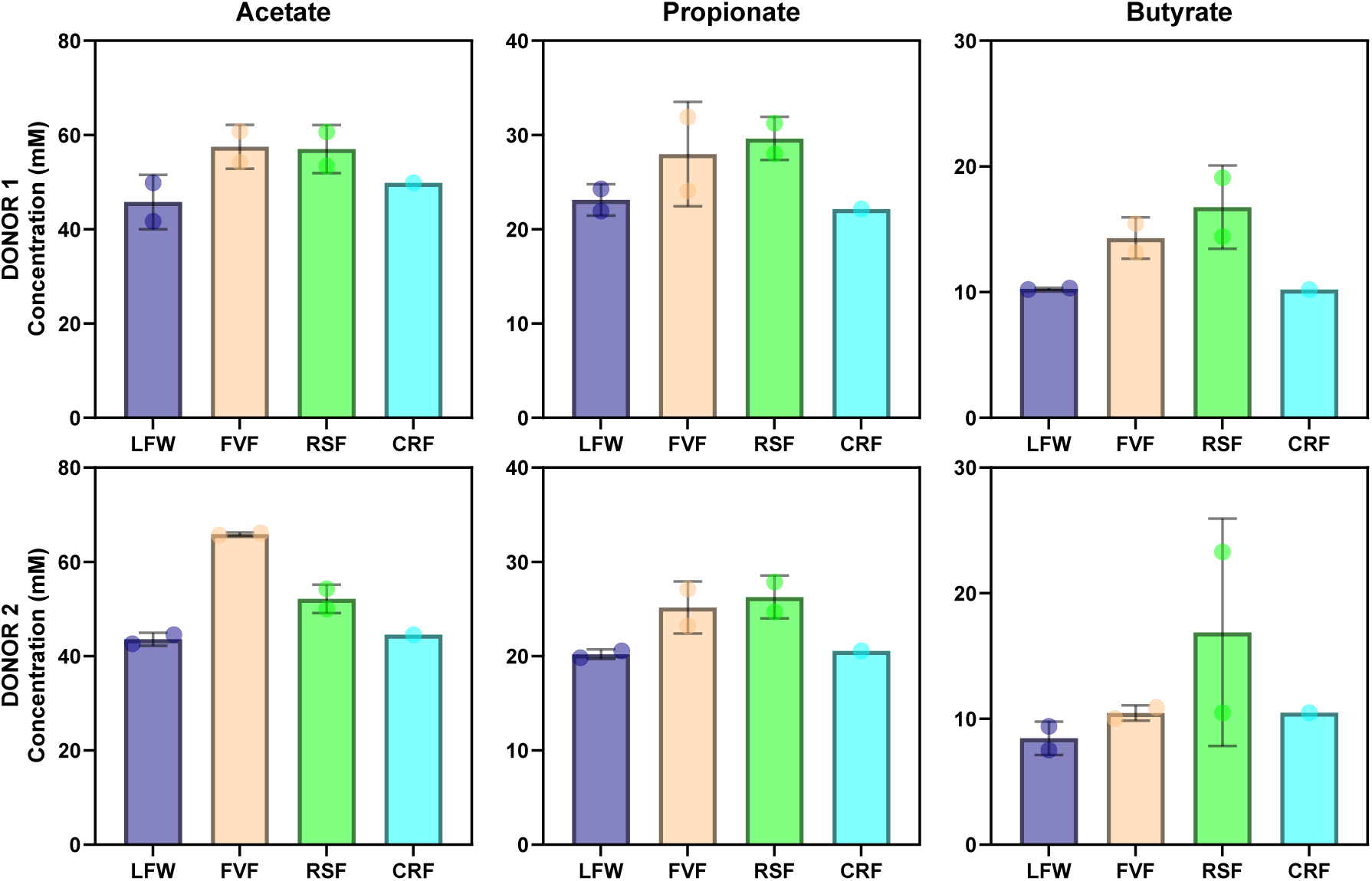
Comparison of Donor 1 and Donor 2 SCFA Concentrations at Day 28 of the Fiber Experiment. In both donors, the supplementation of the FVF and RSF fibers increased all SCFA concentrations, while the CRF fiber did not. The modest increases in SCFA concentrations suggest that the supplementation of the fibers did not significantly affect SCFA concentrations.

**Supplementary Figure 4.**
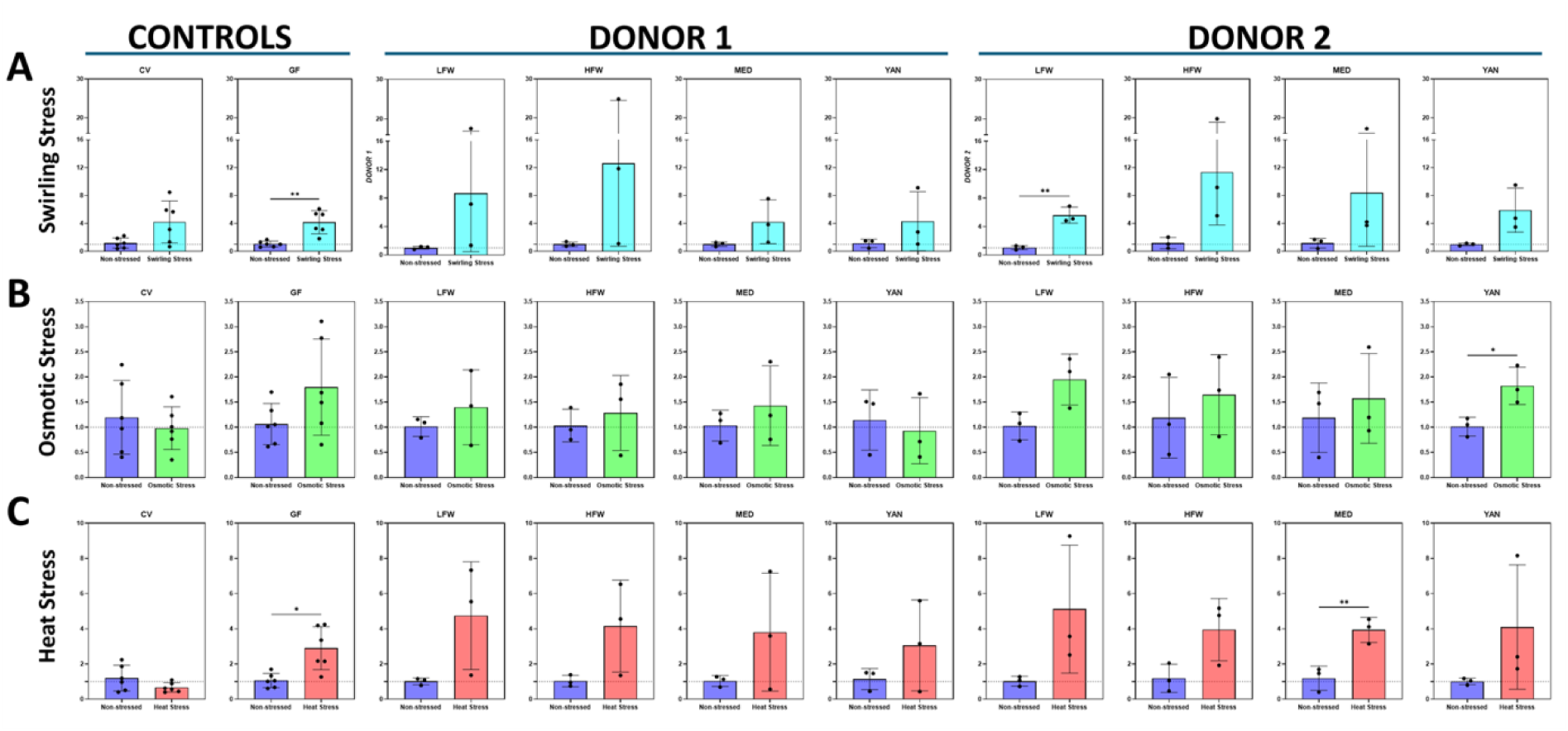
*fosab* Expression in Response to Different Stressors. In the controls, only the swirling stressor increased the expression of *fosab* in the CV group, while all the stressors increased *fosab* expression in the GF group. In majority of the metabolite-treated groups, an overall increase in *fosab* expression was observed although these increases were mostly statistically non-significant.

**Supplementary Figure 5.**
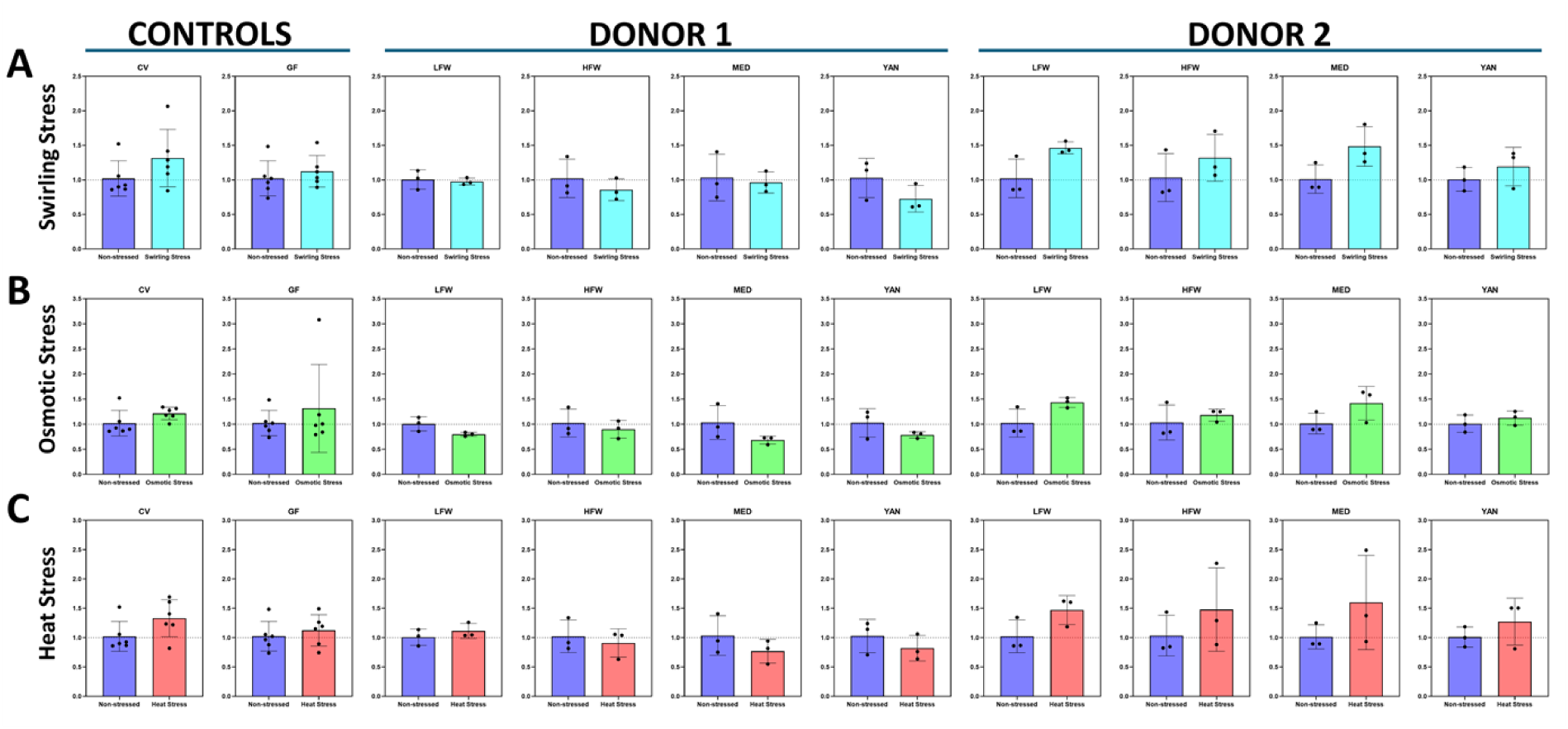
*il-6* Expression in Response to Different Stressors. In the controls, both CV and GF groups had similar response profiles to all the stressors. In zebrafish treated with donor 1 metabolites, *il-6* expression remained relatively similar in non-stressed and stressed conditions. Meanwhile, in zebrafish treated with donor 2 metabolites, *il-6* expression saw modest increases, but none were found to be significant.

**Supplementary Figure 6.**
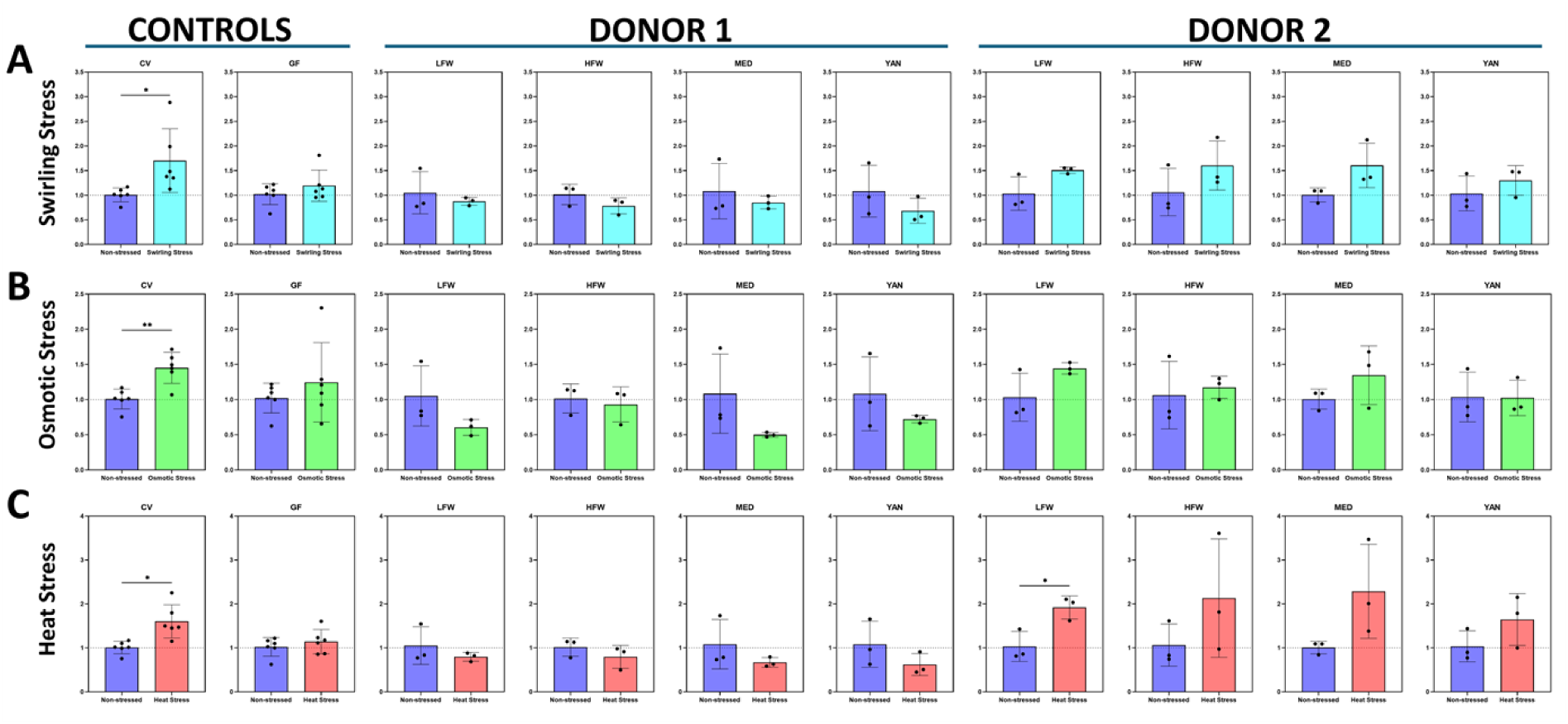
*tnfα* Expression in Response to Different Stressors. In the CV group, all the stressors yielded significant increases in *tnfα* expression, but this response appeared to be blunted in the GF group. In zebrafish treated with donor 1 metabolites, *tnfα* expression appears to be comparable in both non-stressed and stressed conditions. On the other hand, those that were treated with donor 2 metabolites saw moderate increases in *tnfα* expression. Similar trends were observed for *il-6* expression.

